# The complex interaction between megaherbivores, climate and fire has shaped the evolution and distribution of plant spinescence across biogeographical realms

**DOI:** 10.1101/2025.07.02.662719

**Authors:** Rachel Souza Ferreira, Jens J. Ringelberg, Colin Hughes, Eduardo Arlé, Friederike J. R. Wölke, Kyle W. Tomlinson, Renske E. Onstein

**Affiliations:** German Center for Integrative Biodiversity Research (iDiv) Halle – Jena – Leipzig, Puschstrasse 4, 04103 Leipzig, Germany; Leipzig University, Johannisallee 21–23, D-04103 Leipzig, Germany; Naturalis Biodiversity Center, Darwinweg 2, 2333 CR, Leiden, The Netherlands; Biosystematics Group, Wageningen University and Research, Radix Building 107, Droevendaalsesteeg 1, 6708 PB Wageningen, the Netherlands; Department of Systematic and Evolutionary Botany, University of Zurich, Zollikerstrasse 107, CH 8008 Zurich, Switzerland; School of Zoology, George S. Wise Faculty of Life Sciences, Tel Aviv University, Tel Aviv, Israel; Faculty of Environmental Sciences, Czech University of Life Sciences Prague, Kamýcká 129 165 00 Prague, Czech Republic; Center for Integrative Conservation & Yunnan Key Laboratory for Conservation of Tropical Rainforests and Asian Elephants, Xishuangbanna Tropical Botanical Garden, Mengla, Yunnan 666303, China; Center of Conservation Biology, Core Botanical Gardens, Chinese Academy of Sciences, Menglun, Yunnan 666303, China; Leiden University, Institute of Biology Leiden, Sylviusweg 72, 2333 BE Leiden, The Netherlands

**Author notes:** Corresponding Authors: Rachel Souza Ferreira, Renske E. Ontstein.

**Keywords:** anachronism, arms race, defense trait, herbivory, Mimoseae, plant-herbivore interaction

## Abstract

- The evolutionary arms race between plants and herbivores has led to numerous plant adaptations, including spinescence. However, whether spinescence evolved primarily in response to herbivory, or whether abiotic conditions also played a role, remains unknown.
- We integrated phylogenetic, geographic, and trait data for 2,686 species of an ecologically diverse and spiny pantropical lineage – mimosoid legumes – with data for 235 extant and 185 extinct mammalian herbivores >10 kg. Using structural equation models, we assessed how herbivores, climate and fire directly and indirectly affected the proportion of spinescent mimosoids across global and continental assemblages.
- The proportion of spinescent mimosoids in assemblages increased with extant and extinct herbivore richness, drought and heat, while fire influenced spinescence indirectly, by affecting herbivore richness. Notably, correlations of spinescence differed between Africa and America, with increasing importance of extinct herbivores in Africa, illustrating legacy effects on spinescence.
- Megaherbivores have shaped spinescence in mimosoids, especially in dry environments, where losing plant tissues is costly. Spinescence originated ∼35 Mya, with the transition of mimosoids to seasonally dry environments, pre-dating the Miocene expansion of savannas. Our study suggests that both long-term climatic transitions and the emergence of open, herbivore-rich landscapes played crucial roles in the evolution and distribution of spinescence.

## Introduction

The consumption of plant tissues by herbivores has led to an evolutionary arms race between plants and herbivores, leading to the evolution of a diverse range of defense mechanisms and adaptations, expansions in ecological space, and adaptive radiations in both groups (Dawkins, 1979; Endara *et al*., 2017; Perkovich & Ward, 2022). Herbivorous mammals, for example, exhibit diverse feeding behaviours based on the plant parts they consume: grazers primarily consume grasses, while browsers feed on the leaves, flowers and fruits of woody plants (Owen-Smith, 1997). In response, plants have evolved a variety of defense traits to protect themselves from herbivory, including chemical (e.g., highLtannin levels), structural (e.g., spinescence), and symbiotic traits (e.g., bodyguarding ants, associated with *Vachellia* species in Central America and African savannas) (War *et al*., 2012; Palmer, 2023). Among these, structural defenses, such as spines, thorns, or prickles (‘spinescence’ hereafter) are particularly effective against large-bodied herbivores (i.e., herbivores ≥ 10 kg body mass, hereafter ‘megaherbivores’), functioning as barriers that injure the animals’ mouths as they try to feed (Dickison, 2000; Valverde *et al*., 2001)

Although spinescence is generally associated with the presence of herbivores, other factors may also influence its distribution. First, spinescence may be particularly advantageous under certain climatic conditions. For example, in areas with low precipitation and prolonged dry seasons, plants tend to allocate more resources to defense mechanisms, as water scarcity limits photosynthesis and rapid growth, increasing vulnerability to herbivore predation (Grubb, 1992; Vilela *et al*., 2012; Armani *et al*., 2019). This may explain why many spinescent plants are found in seasonally dry, deciduous, fire-intolerant succulent biomes in Madagascar, the Horn of Africa, the Namib, Mexico, Brazil, and Argentina (Ringelberg *et al*., 2020). Second, extinct herbivores may have left an imprint (‘legacy effect’) on the distribution of spinescence in ecosystems today. For example, several areas of the seasonally dry, fire-intolerant succulent biome likely had a historically rich megaherbivore fauna until the Late Pleistocene (ca. 10,000 years ago), but are now relatively poor in mammalian herbivores, with the notable exception of introduced livestock (Mooney *et al*., 1995). Spinescence in these systems may thus represent adaptations to browsing by now extinct megaherbivores, making them evolutionary anachronisms (Janis, 2002).

Nevertheless, spinescent plants remain common in places with high densities of extant (mega-)herbivores, such as fire-prone African savannas (Tomlinson *et al*., 2016; Charles-Dominique *et al*., 2016). Fire and megaherbivores interactively impose disturbance regimes, which maintain open vegetation such as the savannas (Van Langevelde *et al*., 2003). Most woody plant lineages with spinescence in African savannas, such as the ‘Acacias’ (i.e., *Vachellia* and *Senegalia*), diversified from the early Miocene onwards (Greve *et al*., 2012), possibly in response to increasing browsing pressures from megaherbivores (Staver *et al*., 2012; Charles-Dominique *et al*., 2016; Dantas & Pausas, 2020). This diversification also coincided with the rise of bovids and the geographical expansion of African savannas (Beerling & Osborne, 2006; Charles-Dominique *et al*., 2016), creating ecological opportunities for spinescent plants. However, spinescence probably originated much earlier, dating back to the Eocene or even the Cretaceous (e.g., in palms [Arecaceae]) (Onstein *et al*., 2022; Zhang *et al*., 2022).

The drivers of the origin, diversification, and distribution of defense traits and spinescent lineages remain unclear, but spinescence likely resulted from complex interactions between megaherbivores, climate and fire. Notably, Africa and the Americas present striking contrasts in megaherbivore populations, providing a comparative framework to understand the relative importance of historical and contemporary drivers of spinescence (Lehmann *et al*., 2014). Although South America and Africa historically had similar levels of herbivore richness (Owen-Smith, 2013), South America experienced more severe megafauna extinctions during the Pleistocene, losing approximately 80 species (38.65%) of herbivores over 10 kg, while Africa lost only 27 species (17.65%) (Stuart, 2015; Faurby *et al*., 2018). However, declines in distribution ranges are evident in both regions. Such local, global, or functional extinctions of megaherbivores may have substantially changed ecosystems. For example, they could have led to increased woody cover in South American savannas (Doughty *et al*., 2016a) and more frequent wildfires due to the accumulation of plant matter (Galetti, 2004), or even transitions from savanna to forest (Dantas & Pausas, 2022). Past interactions with megafauna may also explain differences in plant traits between Africa and South America, such as more spines (Dantas & Pausas, 2022), higher wood densities, taller plants (Doughty *et al*., 2016b), and larger fruits (Mack, 1993; Wölke *et al*., 2023) in African savannas compared to the South American Cerrado. In contrast, Australia’s extant (mega-)herbivore fauna is now limited to grazing macropods and wombats, and its Late Quaternary extinct assemblage, though including the “giant wombat” (*Diprotodon optatum*), lacked the diverse largeLbodied browsers found in Africa and the Americas (Milewski & Diamond, 2000). Moreover, Australian Pleistocene mammal communities were skewed towards smaller body sizes overall, with proportionately fewer species in the largest size classes than on other continents (Stuart, 2015).

Here, we establish a general comparative framework to study the drivers of the evolution and distribution of spinescence across the global tropics. We use the mimosoid legumes (tribe Mimoseae, formerly subfamily Mimosoideae, Bruneau *et al*., 2024) a pantropical clade of c. 3,500 species of trees, lianas, and shrubs, as a study system. Mimosoids form ecologically important or even dominant elements in all major lowland tropical ecosystems, including rainforests, savannas, and seasonally dry forests (Koenen *et al*., 2020; Ringelberg *et al*., 2023). Importantly, mimosoids have evolved a wide range of defense traits against (mega-)herbivores, including spines, thorns, and prickles, and spinescence evolved independently multiple times throughout mimosoid evolution (Ringelberg *et al*., 2022). This likely facilitated the diversification of mimosoids across the tropics, resulting in radiations of spinescent lineages in different regions, such as *Vachellia* and *Senegalia* in the Americas and African drylands (Greve *et al*., 2012) and mesquites (*Neltuma* and *Strombocarpa*, formerly *Prosopis*) in the seasonally dry tropical scrublands of the succulent biome in the Americas. By contrast, Australian mimosoids (the vast majority of which are in the genus *Acacia*) exhibit a very low proportion of spinescent species, although some taxa have spinescent leaf (phyllode, i.e. modified leaf rachis) tips. We present a newly assembled spinescence trait dataset for 2,686 mimosoid species (76.7% of all species) and combine this with extensive species distribution data and a time-calibrated phylogeny to comprehensively study the evolution and distribution of spinescence across space and time. Given the complex interplay of factors potentially influencing spinescence, our study aims to unravel the interaction between (mega-)herbivores, climate, and fire by testing the following hypotheses:

- **Hypothesis 1**: Species richness of extant and extinct megaherbivores determine global and regional variation in the proportion of spinescent mimosoid species across assemblages, because, to effectively deter herbivory, these structures should have adapted and co-evolved with the feeding behaviour and diversity of herbivores (Burns, 2014; Perkovich & Ward, 2022). Extinct (mega-)herbivores may be particularly important for explaining the distribution of spinescence in regions that lost most megaherbivores in the Pleistocene (i.e., the Americas), reflecting a legacy effect (Stuart, 2015).
- **Hypothesis 2**: Drought, temperature and fire influence spinescence both directly and indirectly, through effects on megaherbivore richness. For example, drought may be directly associated with spinescence due to the increasing vulnerability of plants to herbivory under prolonged dry seasons. However, drought may also correlate with high historical megaherbivore richness, posing an indirect effect on spinescence (e.g., explaining spinescence in fire-intolerant seasonally dry succulent biomes, Ringelberg *et al*., 2020). Similarly, fire frequency may be associated with high extant megaherbivore richness (e.g., explaining spinescence in savanna systems) (Hempson *et al*., 2017) or extinct megaherbivore richness (e.g., in South America, where fire frequency may have increased after the Pleistocene extinctions) (Galetti, 2004), thereby indirectly shaping the distribution of spinescence.

## Material and Methods

### Spatial data resolution and assembly

To test whether the global distribution of spinescence is related to the possible interaction with megaherbivores, we quantified the proportion of spinescent species, species richness of extant and extinct medium-to large-sized mammalian herbivores and average bioclimatic variables and fire frequency across the global distribution of mimosoids. The foundational spatial structure for our analyses was established using an equal area grid cell (Behrmann projection) with a resolution of approximately 100 km (≈1° at the equator). This grid format was utilized throughout our study, when extracting species ranges, bioclimatic variables, and combining these spatial data into an assemblage structure to determine the presence/absence of species in grid cells, or when calculating average climate or fire frequency per grid.

### Mimosoid data

Trait data, focusing on defenses against herbivore predation, were extracted and compiled using the mimosoid species checklist from Ringelberg *et al*. (2023), with generic updates following Bruneau *et al*. (2024). In this classification, the mimosoid clade, formerly known as the subfamily Mimosoideae, comprises around 3,500 species across over 100 genera. Trait information was compiled for 2,686 species spanning all recognized genera. We documented the presence and absence of spinescence (76.7% of species) and spinescence type, defined as prickles (sharp epidermal outgrowths), thorns (woody, modified axillary branches), stipular spines (modified stipules), axillary spines (sharp structures emerging from the leaf axil) or spinescent shoots (shoots transformed into spines) (**Table S1**). To calculate the proportion of plants with spinescence in an assemblage (grid cell), we divided the total number of spinescent species by the sum of all mimosoid species occurring in that grid cell. Occurrence data were adopted from Gagnon *et al*. (2019), who used a curated species checklist to download 444,471 raw records from GBIF, DryFlor, SEINet and other taxon-or region-specific sources. They then updated synonymous names and removed records lacking herbarium vouchers (except DryFlor plot/checklist records), as well as cultivated, marine, centroid or non-native occurrences.

### Environmental data

To test for indirect effects of environmental variables on spinescence, we collected data on annual precipitation (BIO12) and annual mean temperature (BIO1) from CHELSA 1.2 (Karger *et al*., 2017), averaged over one year from 1981 to 2010. Additionally, we calculated the ‘length of the dry season’ as the consecutive months with rainfall < 100 mm/month, using mean monthly precipitation from CHELSA 1.2 (Karger *et al*., 2017). We obtained fire event data from Archibald *et al*. (2013), which used the 0.5° Global Fire Emissions Database 3.1 (GFED) to compile monthly burned area data from 1997 to 2010. We used this dataset to calculate the mean annual proportion of area burned for each grid cell, hereafter referred to as ‘Mean Burn Area’ by applying the ‘join attributes by location’ algorithm in QGIS 3.20.3.

### Extant mammal data

We extracted current polygon ranges of 4,858 extant mammal species from the PHYLACINE database (Faurby *et al*., 2018). ‘Current Ranges’ represent polygon range maps extracted from the International Union for Conservation of Nature, ver. 2016-3 (IUCN, 2016), focusing on areas where species were classified as either extant or native. We defined herbivores as species with a diet with over 95% consisting of plants (i.e., strict herbivores) according to PHYLACINE (Faurby *et al*., 2018). Additionally, we identified grazer and browser species through the MammalDiet database (Kissling *et al*., 2014). For our analysis, we explicitly excluded species that feed exclusively on grasses, also known as ‘exclusive grazers’. Instead, our focus was directed towards species that consume leaves of woody plants, commonly referred to as ‘browsers’ or ‘mixed feeders’, given their potential interaction with mimosoid plants. In addition, we only included species with body mass equal or over 10 kg, following the large mammal definition from Sandom *et al*. (2014). To visualize the overlap between herbivores and mimosoid species, we delineated their ranges using an equal area grid using the *relate* function in the ‘terra’ R package (Hijmans, 2023).

### Extinct mammal data

We derived the probable past distribution of most Late Quaternary extinct medium- and large-bodied terrestrial mammalian herbivore species (353 species) based on the ‘Present Natural’ range in PHYLACINE (Faurby *et al*., 2018). This scenario is based on the distribution of extant species, which, according to fossil records, coexisted with these extinct species. It considers an extinct species to have been present in a particular grid cell if at least 50% of the extant species found in coexistence with it in fossil records were also predicted to occupy that cell before anthropogenic changes. This method is grounded in the assumption that extinct and extant species that shared habitats likely had similar ecological requirements. Therefore, by mapping the pre-human distribution of coexisting extant species, the historical ranges of their extinct counterparts can be inferred (Faurby *et al*., 2018). Within this dataset, we focused on mammals marked with statuses ‘EP’, ‘EX’, and ‘EW’ from IUCN (IUCN, 2016), representing species extinct in prehistory (before 1500 CE), post-1500 CE extinctions, and those extinct in the wild, respectively. Extant species that have become locally extinct are likewise treated as ‘extinct’ in any grid cell where they formerly occurred, so that regional losses of stillLliving taxa are recorded as extinctions on our maps.To explore the potential influence of megaherbivores on plant defense, we included only species with diets comprising at least 95% of plants and body mass equal or over 10 kg, as inferred using PHYLACINE (Faurby *et al*., 2018). However, for extinct species, it is important to note that grazers and browsers cannot be reliably distinguished due to limitations in fossil records and historical data. Therefore, we included all extinct megaherbivores in our analysis.

### Structural Equation Models (SEMs)

To investigate the direct and indirect relationships between the distribution of megaherbivores, climate, fire, and the proportion of spinescent mimosoid species, we used structural equation models (SEMs) implemented in the ‘lavaan’ R package (Rosseel, 2012). We focused on grid cells containing data for more than three mimosoid species, to ensure robust and reliable causal inferences. To assess the sensitivity of this cutoff, we also ran the global SEMs on cells with more than 1 and more than 5 mimosoid species. Our ‘base model’ included direct effects of extant and extinct megaherbivore species richness on the proportion of spinescent mimosoid species, as well as effects of length of the dry season, mean burn area, and mean annual temperature on extant and extinct megaherbivore species richness and the proportion of spinescent mimosoid species (see Table 1 for details on the variables). In addition, we tested whether length of the dry season and mean annual temperature affected mean burn area (i.e., fire frequency). Prior to analysis, mean burn area was log-transformed to meet normality assumptions, and all variables were than scaled to the 0 and 1 range to be able to compare effect sizes between variables and models. We assessed the goodness of fit of the SEMs by verifying that the p-values of χ2-tests were lower than 0.05, the comparative fit index (CFI) exceeded 0.90, and the confidence intervals of the root mean square error of approximation (RMSEA) were below 0.05, following standard practise (Grace *et al*., 2012). We fitted the base model and sequentially removed non-significant model terms until all remaining terms were significant (at P < 0.05). We performed SEMs both globally and within two key biogeographic realms for spinescence distribution in mimosoids: Americas and Africa. We excluded the Caribbean islands, Madagascar, and Réunion due to their distinct histories of megafauna extinctions and associated plant-animal interactions. We did not run models for Australia or Asia, as the proportion of mimosoids with spinescence in these realms is low (but see Discussion).

### Spatial autocorrelation and simulations

We recognize that spatial data might exhibit autocorrelation, given the potential similarity of signals between adjacent grids. To account for the potential impact of spatial autocorrelation on our results, we employed simultaneous autoregressive (SAR) models. SAR models allow for the incorporation of spatial autocorrelation in the residuals of the model. The spatial weights matrix was constructed by identifying the minimum neighbor distance (83.63 km) needed to ensure each cell had at least one adjacent cell, determined via the ‘ncf’ R package (Bjornstad & Cai, 2022) and then calculating weights and Moran’s I values with the ‘spdep’ R package (Bivand *et al*., 2024). We further assessed spatial structure by plotting Moran’s I correlograms for both the raw proportions and the SAR residuals. Standardized coefficients from the SAR models closely matched those from our SEMs (**Tables S2-S4**).

Furthermore, to assess whether the effects of herbivore richness on the global distribution of spinescence could have emerged due to confounding correlations, we performed a sensitivity analysis. Specifically, we repeated the SEM after randomizing the presence/absence of spinescence across mimosoid species 1000 times, maintaining the total mimosoid species richness for each grid cell, and subsequently recalculating the proportion of species with (randomized) spinescence per grid cell. We compared the observed standardized coefficients of extant and extinct megaherbivores on the proportion of spinescence from the empirical SEM to the 1,000 simulated outcomes. We expected that extant and extinct megaherbivores would not account for the distribution of randomized spinescence across assemblages, suggesting that our empirical findings were not influenced by potential confounding correlations between mimosoids, spinescence, and the richness of current and past herbivores.

**Table 1.** Detailed description of variables used in the structural equation models to explain the distribution of spinescence in mimosoid legumes.

| **Variable** | **Description** |
| --- | --- |
| **Proportional richness of spinescent species** | Spinescent species in the grid divided by the total number of species in the grid. |
| **Current megaherbivore richness** | Number of mammalian herbivore species >10kg from the PHYLLACINE database (Faurby *et al.*, 2018). |
| **Extinct megaherbivore richness** | Number of mammalian herbivore species >10kg from the PHYLLACINE database (Faurby *et al.*, 2018). |
| **Length of dry season (drought)** | Consecutive months with rainfall < 100 mm/month from CHELSA, spanning from 1979 to 2010 (Karger *et al.*, 2017). Index from (Ringelberg *et al.*, 2020). |
| **Mean burn area** | Mean annual proportion of area burned for each grid cell (from 1997 to 2010) derived from the pyrome data layer by (Archibald *et al.*, 2013). |
| **Mean annual temperature** | Mean annual temperature from CHELSA, spanning from 1979 to 2013 (Karger *et al.*, 2017). |

### Ancestral state reconstruction of spinescence types

To further evaluate whether spinescence is closely aligned with megaherbivorous mammals and/or abiotic factors, we visually evaluated whether mimosoids evolved concurrently with the diversification of megaherbivores, the spread of African savannas, and/or the expansion of the succulent biome. For this, we conducted ancestral state reconstructions of spinescence types on a time-calibrated mimosoid phylogenetic tree. The mimosoid phylogeny includes 1860 mimosoid species (58% of total mimosoids). The backbone phylogeny was constructed using DNA sequence data obtained through targeted enrichment of 997 nuclear genes (Koenen *et al*., 2020), and combined with 15 species-level phylogenies (Ringelberg *et al*., 2023). We classified mimosoids based on the presence or absence of five topographically non-homologous spinescence types (thorns, axillary spines, prickles, stipular spines, and spinescent shoots), as some species can exhibit multiple types. Species lacking these five types were classified as ‘unarmed’. Species with only spinescent leaf-tips were also considered ‘unarmed’, as leaf-tip spines were not included in our classification, and are only present in few mimosoids. We ran five independent ancestral state reconstructions, one for each spinescence type, and combined the ancestral states subsequently based on the marginal likelihoods of the ancestral spinescence types at the nodes. For each reconstruction, we fit the Equal Rates (ER) and All Rates Different (ARD) models using the *ancr* function in the ‘phytools’ R package (Revell, 2012). The ER model assumes uniform transition rates between states across the phylogeny, whereas the ARD model allows for variable rates between each state transition. To ensure robustness of the findings, each model was repeated 20 times. During each iteration, we used random starting points to avoid local optima and applied log scaling to improve numerical stability. The results from these iterations were evaluated, and the model with the highest log-likelihood was selected as the most likely representation of spinescence evolution. We compared the fit of the ER and ARD models using an analysis of variance (ANOVA), selecting the best model based on the lowest Akaike Information Criterion (AIC) value.

## Results

### Global patterns of spinescent distribution

We reveal distinct distribution patterns among the proportion of spinescent mimosoid species, megaherbivore richness, and environmental factors across global assemblages (**Fig. 1**). Globally, the highest proportions of spinescent mimosoids (**Fig. 1a**) are concentrated in seasonally dry tropical vegetation within the succulent biome (Ringelberg *et al*., 2020), including regions such as northeast Brazil (Caatinga), southwest Madagascar, parts of Mexico, the Horn of Africa, the Karoo–Namib region of southern Africa, and parts of the Gran Chaco in Argentina, Bolivia, and Paraguay. Additionally, while we refer to the Chaco as part of the succulent biome, we acknowledge that its distinction as a separate biome remains debated and therefore treat it as part of the broader seasonally dry vegetation complex. African savannas, particularly in the south and west (e.g., South Africa, Zimbabwe, Mozambique, and the sub-Sahelian zone), also show high spinescence. In contrast, New World savannas, such as the Cerrado, show only modest proportions of spinescent species, levels comparable to those found in the Amazonian rainforest, and much lower than in African savannas. The highest values in the Americas are instead tightly associated with the succulent biome. This contrasts with Africa, where high proportions of spinescence occur in both succulent and savanna biomes. Meanwhile, regions such as the Amazon and Congo basins and most of Australia have low proportions of spinescent mimosoids. Megaherbivore assemblages in the African savanna biome show remarkable extant species richness (**Fig. 1b**), whereas assemblages in Mexico, Brazil, Argentina, and Paraguay show the highest extinct megaherbivore richness (**Fig. 1c**). Globally, the length of the dry season (**Fig. 1e**) is highest in the succulent biome and in the savannas of Australia and parts of southern African savannas, while fire frequency coincides with the distribution of savannas (**Fig. 1d**).

**Figure 1.**
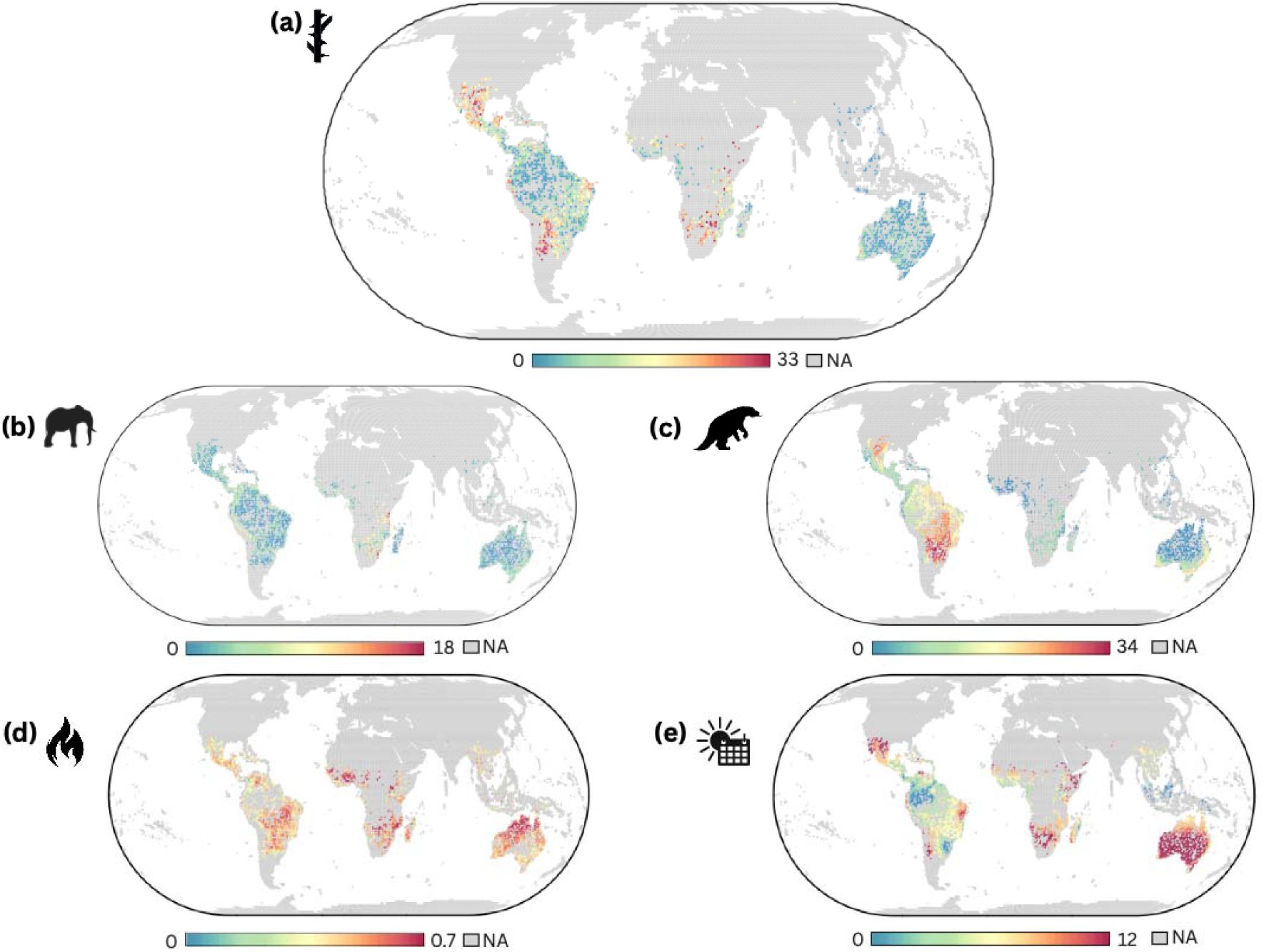
Geographical distribution of spinescence, megaherbivores, and environmental factors across global mimosoid assemblages. Only grid cells (100 x 100 km; ≈1° at the equator) with > 3 mimosoid species are shown. (a) proportion of mimosoid species with spinescence (thorns, axillary spines, prickles, stipular spines or spinescent shoots), (b) extant megaherbivore species richness (herbivorous browsers > 10 kg), (c) extinct megaherbivore species richness (herbivores > 10 kg), (d) mean burn area per year (from 1997 to 2010) (fraction of cell burned per year; values were log-transformed for colour mapping and then back-transformed for the legend) and (e) length of dry season in months (from 1997 to 2010).

### Extant and extinct megaherbivores determine the global distribution of spinescence

The global SEM identified extant megaherbivore richness as the strongest predictor of spinescence (**Fig. 2a, Table S2**), followed by extinct megaherbivore richness, number of dry months and temperature. This indicates that assemblages with greater extant and/or extinct herbivore diversity, with longer dry seasons, and higher temperatures, have a higher proportion of spinescent mimosoid species. Environmental factors also affected spinescence indirectly, via effects on megaherbivore richness. Specifically, we found that extinct and extant megaherbivore richness were higher in cooler areas, and that longer dry seasons were associated with increased extant megaherbivore richness and decreased extinct megaherbivore richness. Furthermore, more frequent fires increased extant megaherbivore richness, suggesting that fire may support herbivore-friendly habitats today. Finally, fire frequency increased in areas with higher mean annual temperatures and longer dry seasons. When using a minimum of 1 or 5 mimosoid species per grid cell, instead of 3, results remained qualitatively similar (**Fig. S1**).

### Continental variation in determinants of spinescence

For the Americas, similar to the global model, the SEM showed the strongest relationship of extant megaherbivore richness on the proportion of spinescent mimosoids (**Fig. 2b, Table S3**), followed by number of dry months and extinct megaherbivore richness. Interestingly, the relationship with temperature suggests that a higher proportion of spinescent mimosoid species is found in cooler rather than warmer places in the Americas, contrasting with global results. The factors affecting extant herbivore richness were largely consistent with the global model. Specifically, extant megaherbivores were found in cooler places with longer dry seasons, but fire frequency decreased rather than increased extant herbivore richness. Contrasting the global model, we found that extinct megaherbivore richness coincided with higher fire frequency. Finally, fire frequency increased with increasing mean annual temperatures.

In Africa, contrasting the global model, the SEM showed the strongest influence of extinct megaherbivore richness on the proportion of spinescent mimosoids (**Fig. 2c, Table S4**), followed by the number of dry months and extant herbivore richness, and no influence of temperature was detected. Similar to the global model, cooler places with longer dry seasons and high fire frequency had higher extant megaherbivore richness, and extinct megaherbivore richness also increased in cooler environments. However, contrasting the global model, extinct herbivore richness also increased with dry season length, and, to a lesser extent, fire. Similar to the Americas, fire frequency increased with increasing annual temperatures.

**Figure 2:**
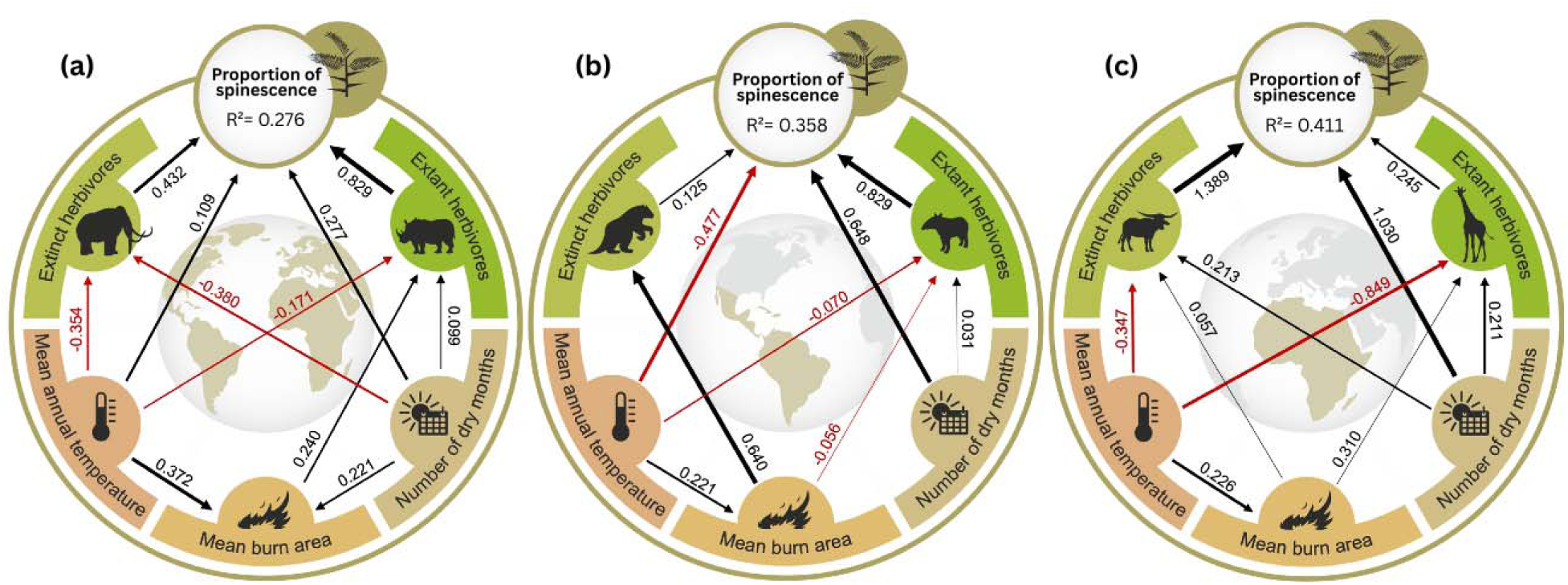
Global and continental determinants of the distribution of spinescent mimosoids across assemblages. Structural equation models showing the direct and indirect effects of extant and extinct megaherbivore species richness and environmental factors on the proportion of spinescent mimosoid species across assemblages with a minimum of 3 mimosoid species. (a) globally (N = 2192 grid cells), (b) in the Americas (N = 1036 grid cells) and (c) in mainland Africa (N = 443 grid cells). ‘Extinct herbivores’ refers to the species richness of extinct [Late Quaternary] megaherbivores, i.e., mammalian herbivores ≥10 kg, ‘Extant herbivores’ to the species richness of mammalian browsing herbivores ≥10 kg, ‘Number of dry months’ to the length of dry season (consecutive months with rainfall < 100 mm/month), ‘Mean burn area’ to fire frequency, and ‘Mean annual temperature’ to the mean annual temperature in degrees Celsius. Boxes represent variables, and arrows indicate the direction of the effects. Arrow thickness is proportional to the strength of the relationships (i.e., standardized path coefficient), with black and red arrows representing significant (p < 0.05) positive and negative path coefficients, respectively. The R² value reflects the total variation explained in the response variable. SEMs across assemblages with a minimum of 1 and 5 mimosoid species can be found in **Fig. S1**.

### Spatial autocorrelation and simulations

The SAR model recovered the same trend of standardized effects as the non-spatial SEM: extant megaherbivore richness exerted the greatest influence on spinescence, followed by the length of the dry season, extinct megaherbivore richness, and mean annual temperature (**Table S2**). Similarly, in the Americas (**Table S3**) and Africa (**Table S4**), SAR coefficients closely matched SEM results. The close correspondence between SAR- and SEM-derived coefficients demonstrates that spatial autocorrelation does not bias the magnitude or ranking of predictor effects.

We further compared the standardized SEM coefficients for each driver of spinescence against 1000 null coefficients obtained by randomizing spinescence across species (while keeping species richness per cell constant). The observed effects of extant and extinct herbivore richness and, to a lesser extent, length of the dry season and mean annual temperature, all exceeded the 95 % interval of their respective null distributions, indicating these relationships are stronger than expected by chance (**Fig. S2**).

### Evolution of spinescence through geological time

The ARD model had the best fit for the evolution of each spinescence type (**Table S5**). The combined ancestral state reconstruction shows that spinescence evolved convergently across mimosoids (**Fig. 3**), with multiple lineages independently evolving spinescence during much of the ca. 40-million-year (Myr) evolutionary history of mimosoid legumes. Furthermore, spinescence likely evolved early, with prickles and stipular spines dating back to at least 35 million years ago (Mya) in some of the older lineages, coinciding with the probable establishment of the succulent biome and substantially predating the expansion of savannas. Based on visual inspections of ancestral state reconstructions, spinescence evolved independently at least 15 times across mimosoid lineages, including in the genera *Mimosa*, *Prosopis*, *Senegalia*, *Vachellia*, and *Acacia*.

**Figure 3.**
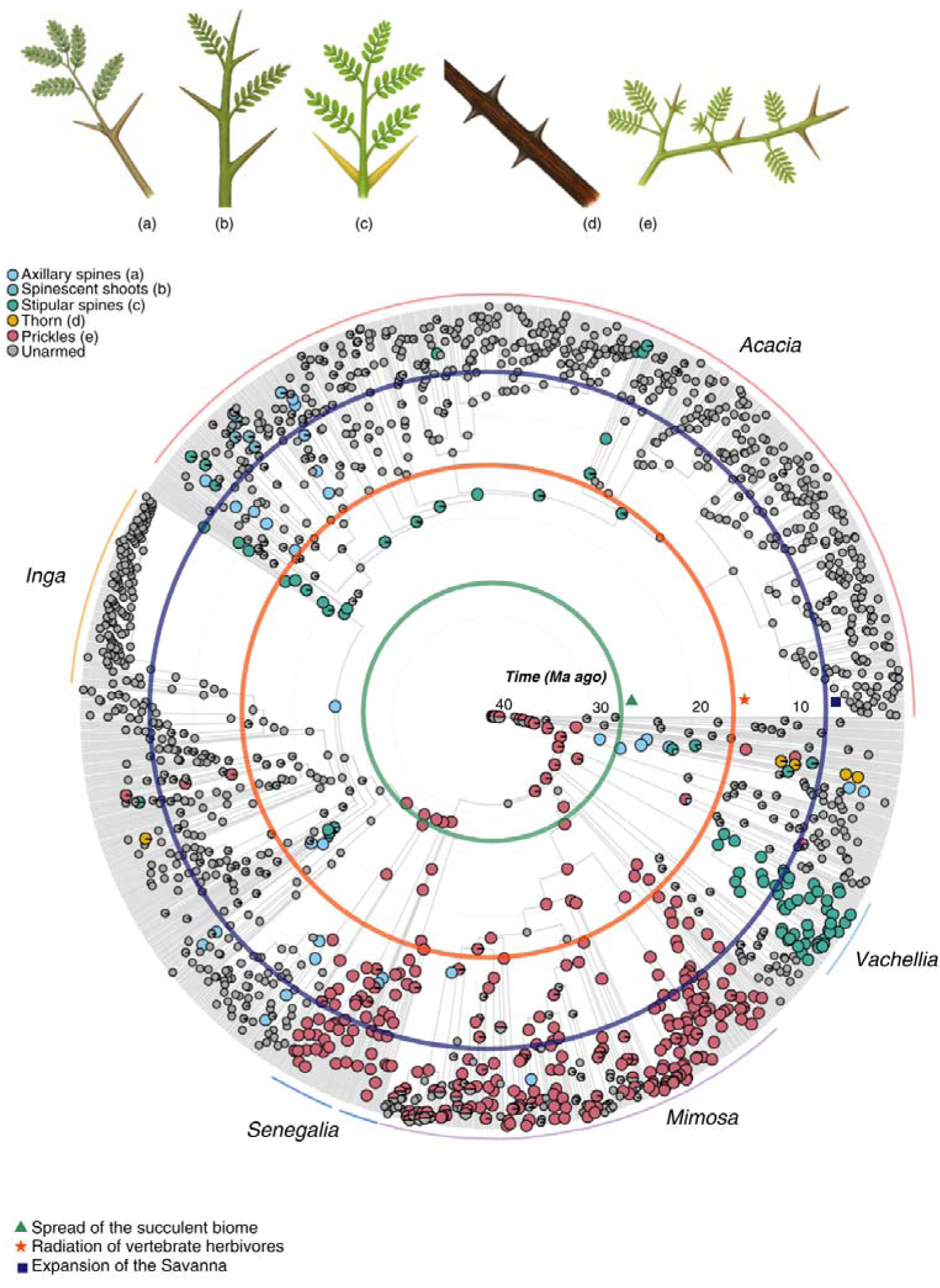
Evolution of spinescence types in mimosoid legumes. Ancestral state reconstructions were performed using maximum likelihood on the mimosoid phylogenetic tree. The pie charts on nodes represent the probability of the five different spinescence types and the absence of spinescence (‘unarmed’) under the best fitting models (i.e., All Rates Different, ARD; **Table S5**). Prominent mimosoid genera are highlighted at the tips of the phylogenetic tree. Biotic and environmental events are represented with symbols and rings placed at their period of occurrence, with probable establishment of the succulent biome at ca. 34 – 23 Mya (Gagnon *et al*., 2019), radiations of mammalian mega-herbivores (e.g., bovids) in the mid-Miocene at ca. 15 Mya (Charles-Dominique *et al*., 2016), and the expansion of savannas starting at ca. 15 – 8 Mya (Cerling *et al*., 1997).

## Discussion

We explored the relationship between the distribution of spinescence and megaherbivore richness, alongside environmental factors such as temperature, fire regimes, and drought (**Fig. 1**). Using the mimosoid clade as a model system, our findings strongly support our hypothesis (**H1**) that extant and extinct megaherbivores are key predictors of spinescence across global plant assemblages (**Fig. 2**). However, continental analyses reveal that extinct megaherbivore richness was a particularly strong predictor of spinescence in Africa, but not in the Americas, contrasting our expectation. Furthermore, we show that drought, temperature and fire are also crucial for explaining the distribution of spinescence, either directly, or indirectly, via effects on extant and extinct megaherbivore richness – supporting our second hypothesis (**H2**). Specifically, we found that drought consistently increases the proportion of spines in assemblages, illustrating how physical defence traits are especially important in seasonally dry environments, where losing plant parts due to herbivory is likely to be particularly costly. Deciduousness and the need to invest in rapid leaf flush and growth at the start of the wet season each year is ubiquitous across woody species, including mimosoids, in the succulent biome, further illustrating the importance of plant defence in dry environments. Length of the dry season is also associated with high megaherbivore richness historically in Africa, consistent with the high frequency of spinescence in the succulent biome. Furthermore, it is notable that the first appearance of spinescence in mimosoids coincided with an evolutionary shift from taxa confined to tropical wet forests to lineages in seasonally dry tropical climates (Ringelberg *et al*., 2023), suggesting that drought and the global expansion of aridity was an important driver of the evolution of spinescence well before expansion of the savanna biome in the Miocene. Fire enhances extant megaherbivore richness globally and in Africa, explaining spinescence in savanna systems, and had a strong association with extinct megaherbivore richness in the Americas, consistent with the increase in fire frequency after the Pleistocene extinctions (Galetti, 2004). Finally, we demonstrate that spinescence arose independently at different times across mimosoid lineages, indicating convergent evolution of their defense structures (**Fig. 3**). This provides further evidence that both biotic and abiotic selection pressures have synergistically shaped spinescence in mimosoid legumes.

### Extant herbivores influence the broad-scale distribution of spinescence

Our structural equation models highlight both the differences and similarities in the factors influencing the proportion of spinescent mimosoids globally, in the Americas, and on mainland Africa. Globally and in the Americas, the richness of extant mammalian herbivores emerged as the primary driver of spinescence (**Fig. 2a, b**). In Africa, extant herbivore richness also influenced the proportion of spinescence, but with a weaker effect (**Fig. 2c**). This finding aligns with the evolutionary arms race theory (Dawkins & Krebs, 1979) and the distribution of defense traits against browsing pressure, e.g., in savanna mimosoid genera (Greve *et al*., 2012). Our findings reveal a striking result that highlights the strong effect of extant herbivores on spinescence in South America, even though most of its megafauna has gone extinct. This suggests that the current browsing pressure from herbivores may still play a significant role in shaping plant defenses. Similar evidence was reported for South American palms, where tapirs, deer and peccaries had strong influence on the presence of defense traits

### Extinct megaherbivores and their ecological legacy

While the primary distributional patterns of extinct megaherbivores are robustly supported by extensive fossil evidence, museum specimens, and the scientific literature, enabling the assembly of a comprehensive dataset (Faurby *et al*., 2018), the geographic scope of these patterns should be interpreted with caution, since reconstructed ranges remain vulnerable to spatial gaps arising from taxonomic sampling biases and uneven fossil preservation (Holland, 2016). Although we recognize these limitations, we demonstrate that extinct megaherbivores have nonetheless left lasting ecological imprints on modern ecosystems, especially in the abundance of spinescent plants, as predicted by our hypotheses and reveled by our spatial analyses (**Fig. 2**). In South America, for example, the diversity of extinct large browsers (Owen-Smith, 2013) suggests that woody plants were subject to intense herbivory pressure, leading to the evolution of mechanical defenses such as spines and thorns. In several mimosoid species, such as *Mimosa guanacastensis*, *Pithecellobium platylobum*, and *Senegalia riparia*, spinescence may have originally evolved as deterrents to large extinct herbivores such as ground sloths and gomphotheres (Janzen & Martins, 1982). While these defenses may now be subject to reduced selection pressures, their persistence highlights the lingering impact of past herbivory. Similarly, megafaunal fruits in Neotropical palm species may be relicts of dispersal by now extinct Pre-Quaternary megafauna (Lim *et al*., 2020; Wölke *et al*., 2023). In Africa, our results indicate that locally or globally extinct megaherbivores are the major factor influencing the proportion of spinescence. This pattern aligns with findings in African savannas, where plants with spines and densely branched crowns may have evolved in response to now extinct megaherbivores (Staver et al., 2012; Charles-Dominique et al., 2016; Anest et al., 2024), highlighting how the legacy of extinct herbivores has influenced the distribution and evolution of spinescence across diverse biogeographical realms.

### Complex ecosystem dynamics: The role of climate and fire

While megaherbivores play a central role in shaping plant defenses, environmental factors such as climate and fire regimes also influence spinescence. In our study, the length of the dry season directly impacted the proportion of spinescent species globally, in the Americas and in Africa, while fire frequency had an indirect but notable influence by affecting megaherbivore richness globally and in Africa. Spinescent plants are commonly found in open, arid, and seasonally dry environments, such as savannas, where herbivory pressure is high and water availability is low (Charles-Dominique *et al*., 2016; Dantas & Pausas, 2022). However, our results show that spinescence is not exclusive to savannas (**Fig. 1**). Across the global succulent biome, which is characterized by frost-free climates, prolonged drought, abundance of fire-intolerant stem succulents, deciduousness, and relative absence of C4 grasses, spinescence is also prevalent (Gagnon *et al*., 2019; Ringelberg *et al*., 2020). Indeed, it is striking that in the Neotropics, areas with the highest proportions of spinescent mimosoids coincide closely with the distribution of the succulent biome *sensu* Ringelberg *et al*. (2020). This pattern likely reflects the fact that in dry environments, where plant growth and tissue replacement are slow and deciduousness is almost universal, the impact of herbivory is more severe. Under these conditions, defenses such as spines confer a critical advantage by deterring browsers before damage occurs (Coley *et al*., 1985; Herms & Mattson, 1992). During the Pliocene, increasing drought and the establishment of open grasslands were tightly coupled with rises in megaherbivore richness (Bobe, 2006), demonstrating a historical association between aridity and megafauna diversity in seasonally dry ecosystems in Africa. Present day succulent biomes are also rich in megaherbivores, especially ungulates, including the grey rehbock (*pelea capreolus*) and bontebok (*Damaliscus pygargus*) (Radloff, 2008). This illustrates how spines in (seasonally-)dry places, e.g., succulent biomes, may reflect both current and legacy effects of historical herbivory.

The positive correlation between fire frequency, extant megaherbivore richness, and spinescence likely reflects the dynamic balance between fire and herbivory over time. This balance may drive dynamic shifts in the distribution of fire-tolerant versus browse-tolerant trees in a savanna (Bond *et al*., 2001). Our results reflect these complex dynamics by showing that fire frequency, temperature and dry season length significantly impact megaherbivore distributions and spinescence in Africa (**Fig. 2c**). In the Americas, our findings indicate that regions with extended dry periods and slightly lower temperatures support higher megaherbivore richness. Conversely, Neotropical savannas with high fire frequency tend to have lower megaherbivore richness, but higher past megaherbivore diversity, illustrating how fire, together with the ecological legacy of extinct megaherbivores, has served as a joint ecosystem modifier (Galetti, 2004; Bond & Keeley, 2005). Interestingly, the proportion of spinescent mimosoids in the Cerrado is low compared to other seasonally dry Neotropical vegetation, and similar to proportions in e.g., the Amazon rainforest (**Fig. 1a**). Even though the Cerrado historically supported a rich diversity of megaherbivores (Paulo & Bertini, 2016), its acidic, nutrient-poor soils may have resulted in weaker selection pressure for spinescence. In response, Cerrado woody plants appear to have redirected defensive investments toward tougher wood, potent chemical compounds, and alternative deterrents such as latex secretion or spines on fruits (Dantas & Pausas, 2022). This shift in defense strategy likely explains the low spinescence richness observed despite high past megafauna richness in the region.

Our findings also highlight the low proportion of spinescence in certain regions and biomes (**Fig. 1**). In rainforests, plants often rely on chemical defenses to deter primarily insect herbivores (Mithöfer & Boland, 2012). Interestingly, while spinescence tends to be more common in dry areas, it is notably less prevalent in Australia (**Fig. 1**). This pattern is likely due to the predominance of the large (> 1,000 species) mostly non-spinescent mimosoid genus *Acacia* in Australia, which comprises mainly evergreen species with reduced phyllodinous leaves. In some cases, the phyllodes can be hardened and pointed, potentially serving a defensive role. Gélin *et al*., (2023), who surveyed spine origins across a diverse array of eudicot lineages, and Nge *et al*., (2024), who focused on spine evolution in the genus *Cryptandra* (Rhamnaceae), both reported Pliocene radiations of spinescent taxa in Australia, suggesting a broader pattern of delayed spine evolution on that continent (**Fig. 3**).

### The evolution of spinescence

Although our spatial analyses provide strong evidence for the interactive effects of herbivores, climate and fire on the distribution of spinescence, ancestral state reconstructions illustrate a more complicated evolutionary history of spinescence types (**Fig. 3**). Different types, such as prickles or thorns, which are not homologous, emerged at different times, some (e.g., prickles and stipular spines) possibly dating back to ca. 35 Mya. This early timing is corroborated by Zhang *et al*. (2022) who recovered a diverse assemblage of spiny-plant fossils of prickles and thorns from late Eocene (∼39 Mya) contemporaneous with the emergence of open semi-wooded vegetation in central Tibet. This timing also coincides with the shift of mimosoids to seasonally-dry succulent biomes (Gagnon *et al*., 2019; Ringelberg *et al*., 2023), and the (re-)emergence of megaherbivores after the ‘megaherbivore gap’ at ca. 40 Mya (Smith *et al*., 2010; Onstein *et al*., 2022), but it substantially predates the Miocene expansion of savannas (Arakaki *et al*., 2011; Pennington & Hughes, 2014). Nevertheless, we observe a notable diversification of spinescent lineages beginning in the mid- to late Miocene (from ca. 15 Mya onwards, **Fig. 3**), consistent with the spread of savanna ecosystems and the diversification of bovids (Beerling & Osborne, 2006; Charles-Dominique *et al*., 2016). We suggest that trait- and climate-dependent diversification rate models can be used to more quantitatively test the association between historical climate, megaherbivores, defence evolution and diversification in mimosoids and other spinescent angiosperms (e.g., Anest *et al.,* 2024).

## Acknowledgements

RSF and REO acknowledge support from iDiv, funded by the German Research Foundation (DFG FZT 118, 202548816). We thank members of iDiv’s Evolution and Adaptation research group and Mauro Galetti for their valuable insights, and Gwilym P. Lewis for his assistance during data collection. We are grateful to Sally Archibald for kindly providing the fire history data used in our analyses. We thank Søren Faurby for a helpful suggestion after a conference talk in Prague that helped shape the direction of our analysis. EA thanks the funding provided by the Azrieli Foundation through their International Postdoctoral Fellowship Programme, in association with the Belmaker Lab – Marine Ecology and Biodiversity. We are also grateful to Kew Gardens for providing access to the specimens in the herbarium. Additionally, we acknowledge the Helmholtz Centre for Environmental Research for providing access to EVE, a High-Performance Computer.

## Data availability

All data needed to evaluate the conclusions in the paper are present in the paper and/or the Supplementary Materials. R scripts used for all analyses in this study will be made available in the Dryad Digital Repository upon publication in a peer-reviewed journal. Occurrence datasets and the mammal and mimosoid trait datasets have been deposited on Zenodo available for reviewers using the following link: https://zenodo.org/records/15773619?token=eyJhbGciOiJIUzUxMiJ9.eyJpZCI6IjNmNGZkNDE0LTY0N2MtNDBlYS04YmYyLTg5ZGQ3N2YxZDIwMCIsImRhdGEiOnt9LCJyYW5kb20iOiI1ODQ0NTIzODBhYmM0MTIzMzViODgzODA4MTY3MjZhYyJ9.IXMrzZWk8h50ktSZtIhczUDJbkgaDA-VXGmEyofULE2t8giGeGlEXAGX8WOuAd_MqvHKmwAvAQxzasN3RSxZ1Q

## Author contributions

REO and RSF designed the study, with input from JJR and CH. RSF collected the data and ran analyses with contributions from REO, JJR, CH, FJRW and EA. RSF and REO wrote the manuscript with contributions from JJR, CH, EA, FJRW and KT.

## References

Anest, A., Bouchenak-Khelladi, Y., Charles-Dominique, T., Forest, F., Caraglio, Y., Hempson, G. P., Maurin, O. & Tomlinson, K. W. (2024). Blocking then stinging as a case of two-step evolution of defensive cage architectures in herbivore-driven ecosystems. Nature Plants, 10(4), pp. 587–597. doi:10.1038/s41477-024-01649-4

Arakaki, M., Christin, P., Nyffeler, R., Lendel, A., Eggli, U., Ogburn, R.M., Spriggs, E., Moore, M.J. & Edwards, E.J. (2011). Contemporaneous and recent radiations of the world’s major succulent plant lineages, Proceedings of the National Academy of Sciences of the United States of America 108(20), pp. 8379–8384. doi:10.1073/pnas.1100628108.

Archibald, S., Lehmann, C. E. R., Gómez-Dans, J. L. & Bradstock, R. A. (2013). Defining pyromes and global syndromes of fire regimes. Proceedings of the National Academy of Sciences of the United States of America, 110, pp. 6442–6447. doi:10.1073/pnas.1211466110

Armani, M., Charles-Dominique, T., Barton, K. E. & Tomlinson, K. W. (2019). Developmental constraints and resource environment shape early emergence and investment in spines in saplings. Annals of Botany, 124(7), pp. 1133–1142. doi:10.1093/aob/mcz152

Beerling, D. J. & Osborne, C. P. (2006). The origin of the savanna biome. Global Change Biology, 12, pp. 2023–2031. doi:10.1111/j.1365-2486.2006.01239.x

Bjornstad, O. N. & Cai, J. (2022). *ncf*: spatial covariance functions (R package version 1.3–2). Available at: https://cran.r-project.org/web/packages/ncf/ncf.pdf.

Bivand, R., Altman, M., Anselin, L., Assunção, R., Bera, A. et al. (2024). *spdep*: spatial dependence: weighting schemes, statistics (R package version 1.3–11). Available at: https://cran.r-project.org/web/packages/spdep/spdep.pdf.

Bobe, R. (2006). The evolution of arid ecosystems in eastern Africa. Journal of Arid Environments, 66(3), pp. 564–584. doi:10.1016/j.jaridenv.2006.01.010

Bond, W. J. & Keeley, J. E. (2005). Fire as a global ‘herbivore’: the ecology and evolution of flammable ecosystems. Trends in Ecology & Evolution, 20, pp. 387–394. doi:10.1016/j.tree.2005.04.025

Bond, W. J., Smythe, K. A. & Balfour, D. A. (2001). Acacia species turnover in space and time in an African savanna. Journal of Biogeography, 28, pp. 117–128. doi:10.1046/j.1365-2699.2001.00506.x

Bruneau, A., de Queiroz, L. P., Ringelberg, J. J., Borges, L. M., da Costa Bortoluzzi, R. L., Brown, G. K., Cardoso, D. B., Clark, R. P., de Souza Conceição, A., Cota, M. M. T. & Demeulenaere, E. (2024). Advances in Legume Systematics 14. Classification of Caesalpinioideae. Part 2: Higher-level classification. PhytoKeys, 240, pp. 1–552. doi:10.3897/phytokeys.240.101716

Burns, K. C. (2014). Are there general patterns in plant defence against megaherbivores? Biological Journal of the Linnean Society, 111, pp. 38–48. doi:10.1111/bij.12181

Cerling, T. E., Harris, J. M., MacFadden, B. J., Leakey, M. G., Quade, J., Eisenmann, V. & Ehleringer, J. R. (1997). Global vegetation change through the Miocene/Pliocene boundary. Nature, 389(6647), pp. 153–158. doi:10.1038/38229

Charles-Dominique, T., Davies, T. J., Hempson, G. P., Bezeng, B. S., Daru, B. H., Kabongo, R. M., Maurin, O., Muasya, A. M., Van der Bank, M. & Bond, W. J. (2016). Spiny plants, mammal browsers, and the origin of African savannas. Proceedings of the National Academy of Sciences of the United States of America, 113(38), pp. E5572–E5579. doi:10.1073/pnas.1607493113

Coley, P. D., Bryant, J. P. & Chapin, F. S. III. (1985). Resource availability and plant antiherbivore defense. Science, 230(4728), pp. 895–899. doi:10.1126/science.230.4728.895

Dantas, V. L. & Pausas, J. G. (2020). Megafauna biogeography explains plant functional trait variability in the tropics. Global Ecology and Biogeography, 29, pp. 1288–1298. doi:10.1111/geb.13111

Dantas, V. L. & Pausas, J. G. (2022). The legacy of the extinct Neotropical megafauna on plants and biomes. Nature Communications, 13, pp. 129. doi:10.1038/s41467-021-27749-9

Dawkins, R. & Krebs, J. R. (1979). Arms races between and within species. Proceedings of the Royal Society B, 205, pp. 489–511. doi:10.1098/rspb.1979.0081

Dickison, W. C. (2000). Integrative Plant Anatomy. San Diego, CA, USA: Academic Press.

Doughty, C. E., Faurby, S. & Svenning, J.-C. (2016a). The impact of the megafauna extinctions on savanna woody cover in South America. Ecography, 39, pp. 213–222. doi:10.1111/ecog.01593

Doughty, C. E., Wolf, A., Malhi, Y. & Svenning, J.-C. (2016b). Megafauna extinction, tree species range reduction, and carbon storage in Amazonian forests. Ecography, 39, pp. 194–203. doi:10.1111/ecog.01715

Endara, M. J., Coley, P. D., Ghabash, G., Nicholls, J. A., Dexter, K. G., Donoso, D. A., Stone, G. N., Pennington, R. T. & Kursar, T. A. (2017). Coevolutionary arms race versus host defense chase in a tropical herbivore–plant system. Proceedings of the National Academy of Sciences of the United States of America, 114(36), pp. E7499–E7505. doi:10.1073/pnas.1707727114

Faurby, S., Davis, M., Pedersen, R. Ø., Schowanek, S. D., Antonelli, A. & Svenning, J.-C. (2018). PHYLACINE 1.2: the Phylogenetic Atlas of Mammal Macroecology. Ecology, 99, pp. 2626. doi:10.1002/ecy.2443

Galetti, M. (2004). Parks of the Pleistocene: recreating the Cerrado and the Pantanal with megafauna. Natureza e Conservação, 2, pp. 93–100.

Galetti, M., Moleón, M., Jordano, P., Pires, M. M., Guimarães Jr, P. R., Pape, T., Svenning, J.-C. et al. (2018). Ecological and evolutionary legacy of megafauna extinctions. Biological Reviews, 93(2), pp. 845–862. doi:10.1111/brv.12374

Gélin, U., Charles-Dominique, T., Davies, T. J., Svenning, J.-C., Bond, W. J. & Tomlinson, K. W. (2023). The evolutionary history of spines – a Cenozoic arms race with mammals. bioRxiv [preprint]. doi:10.1101/2023.02.09.527903

Gagnon, E., Ringelberg, J. J., Bruneau, A., Lewis, G. P. & Hughes, C. E. (2019). Global succulent biome phylogenetic conservatism across the pantropical *Caesalpinia* group (Leguminosae). New Phytologist, 222, pp. 1994–2008. doi:10.1111/nph.15819

Grace, J. B., Schoolmaster Jr, D. R., Guntenspergen, G. R., Little, A. M., Mitchell, B. R., Miller, K. M. & Schweiger, E. W. (2012). Guidelines for a graph-theoretic implementation of structural equation modeling. Ecosphere, 3(8), pp. 1–44. doi:10.1890/ES12-00048.1

Greve, M., Lykke, A. M., Fagg, C. W., Bogaert, J., Friis, I., Marchant, R., Marshall, A. R., Ndayishimiye, J., Sandel, B. S., Sandom, C. & Schmidt, M. (2012). Continental-scale variability in browser diversity is a major driver of diversity patterns in acacias across Africa. Journal of Ecology, 100(5), pp. 1093–1104. doi:10.1111/j.1365-2745.2012.01994.x

Göldel, B., Araujo, A. C., Kissling, W. D. & Svenning, J.-C. (2016). Impacts of large herbivores on spinescence and abundance of palms in the Pantanal, Brazil. Botanical Journal of the Linnean Society, 182, pp. 465–479. doi:10.1111/boj.12421

Grubb, P. J. (1992). Positive distrust in simplicity: lessons from plant defences and from competition among plants and among animals. Journal of Ecology, 80, pp. 585–610. doi:10.2307/2260897

Hempson, G. P., Archibald, S. & Bond, W. J. (2017). The consequences of replacing wildlife with livestock in Africa. Scientific Reports, 7, pp. 17196. doi:10.1038/s41598-017-17250-4

Herms, D. A. & Mattson, W. J. (1992). The dilemma of plants: to grow or defend. The Quarterly Review of Biology, 67(3), pp. 283–335. doi:10.1086/417895

Hijmans, R. J. (2023). terra: spatial data analysis (R package version 1.8–4). Available at: https://CRAN.R-project.org/package=terra.

Holland, S. M. (2016). The non-uniformity of fossil preservation. Philosophical Transactions of the Royal Society B: Biological Sciences, 371(1699), pp. 20150130. doi:10.1098/rstb.2015.0130

Irlbeck, N. A. & Hume, I. D. (2003). The role of *Acacia* in the diets of Australian marsupials? A review. Australian Mammalogy, 25(2), pp. 121–134. doi:10.1071/AM03014

IUCN. (2016). The IUCN Red List of Threatened Species, Version 2025-1. Available at: https://www.iucnredlist.org. Downloaded on 13 June 2022.

Janis, C. M. (2002). The ghosts of evolution: nonsensical fruit, missing partners and other ecological anachronisms. Annual Review of Ecology and Systematics, 17, pp. 595–636. doi:10.1146/annurev.ecolsys.17.1.595

Janzen, D. H. & Martin, P. S. (1982). Neotropical anachronisms: the fruits the gomphotheres ate. Science, 215, pp. 19–27. doi:10.1126/science.215.4528.19

Karger, D. N., Conrad, O., Böhner, J., Kawohl, T., Kreft, H., Soria-Auza, R. W., Zimmermann, N. E., Linder, H. P. & Kessler, M. (2017). Climatologies at high resolution for the earth’s land surface areas. Scientific Data, 4, pp. 170122. doi:10.1038/sdata.2017.122

Kissling, W. D., Dalby, L., Fløjgaard, C., Lenoir, J., Sandel, B., Sandom, C., Trøjelsgaard, K. & Svenning, J.-C. (2014). Establishing macroecological trait datasets: digitalization, extrapolation, and validation of diet preferences in terrestrial mammals worldwide. Ecology and Evolution, 4(14), pp. 2913–2930. doi:10.1002/ece3.1125

Koenen, E. J. M., Kidner, C., de Souza, É. R., Simon, M. F., Iganci, J. R., Nicholls, J. A., Brown, G. K., de Queiroz, L. P., Luckow, M., Lewis, G. P., Pennington, R. T. & Hughes, C. E. (2020). Hybrid capture of 964 nuclear genes resolves evolutionary relationships in the mimosoid legumes and reveals the polytomous origins of a large pantropical radiation. American Journal of Botany, 107(12), pp. 1710–1735. doi:10.1002/ajb2.1568

Lehmann, C. E. R., Archibald, S., Hoffmann, W. A. & Bond, W. J. (2014). Savanna vegetation– fire–climate relationships differ among continents. Science, 343, pp. 548–552. doi:10.1126/science.1247356

Mack, A. L. (1993). The sizes of vertebrate-dispersed fruits: a neotropical-paleotropical comparison. The American Naturalist, 142, pp. 840–856. doi:10.1086/285571

Milewski, A. V. & Diamond, R. E. (2000). Why are very large herbivores absent from Australia? A new theory of micronutrients. Journal of Biogeography, 27(4), pp. 957–978. doi:10.1046/j.1365-2699.2000.00436.x

Mithöfer, A. & Boland, W. (2012). Plant defense against herbivores: chemical aspects. Annual Review of Plant Biology, 63, pp. 431–450. doi:10.1146/annurev-arplant-042110-103854

Mooney, H. A., Bullock, S. H. & Medina, E. (1995). Introduction. In: Bullock, S. H., Mooney, H. A. & Medina, E. (eds), Seasonally dry tropical forests, pp. 1–8. Cambridge, UK: Cambridge University Press.

Nge, F. J., Kellermann, J., Biffin, E., Thiele, K. R. & Waycott, M. (2024). Rise and fall of a continental mesic radiation in Australia: spine evolution, biogeography and diversification of *Cryptandra* (Rhamnaceae: Pomaderreae). Botanical Journal of the Linnean Society, 204, pp. 327–342. doi:10.1093/botlinnean/boad051

Onstein, R. E., Kissling, W. D. & Linder, H. P. (2022). The megaherbivore gap after the non-avian dinosaur extinctions modified trait evolution and diversification of tropical palms. Proceedings of the Royal Society B, 289, pp. 20212633. doi:10.1098/rspb.2021.2633

Owen-Smith, N. (1997). Distinctive features of the nutritional ecology of browsing versus grazing ruminants. Zeitschrift für Säugetierkunde, 62, pp. 176–190.

Owen-Smith, N. (2013). Contrasts in the large herbivore faunas of the southern continents in the late Pleistocene and the ecological implications for human origins. Journal of Biogeography, 40, pp. 1215–1224. doi:10.1111/jbi.12100

Palmer, T. M. (2023). Acacia ants. Current Biology, 33(11), pp. R469–R471. doi:10.1016.j.cub.2023.02.002

Paulo, P. O., & Bertini, R. J. (2015). Mamíferos fósseis do limite Pleistoceno/Holoceno do estado de Goiás. Revista do Instituto Geológico (Descontinuada), 36(2), pp. 61–75. doi:10.5935/0100-929X.20150008

Perkovich, C. & Ward, D. (2022). Differentiated plant defense strategies: herbivore community dynamics affect plant–herbivore interactions. Ecosphere, 13, pp. e3935. doi:10.1002/ecs2.3935

Revell, L. J. (2024). phytools 2.0: an updated R ecosystem for phylogenetic comparative methods (and other things). PeerJ, 12, pp. e16505. doi:10.7717/peerj.16505

Ringelberg, J. J., Zimmermann, N. E., Weeks, A., Lavin, M. & Hughes, C. E. (2020). Biomes as evolutionary arenas: convergence and conservatism in the trans-continental succulent biome. Global Ecology and Biogeography, 29(7), pp. 1100–1111. doi:10.1111/geb.13089

Ringelberg, J. J., Koenen, E. J., Iganci, J. R., de Queiroz, L. P., Murphy, D. J., Gaudeul, M., Bruneau, A., Luckow, M., Lewis, G. P. & Hughes, C. E. (2022). Phylogenomic analysis of 997 nuclear genes reveals the need for extensive generic re-delimitation in Caesalpinioideae (Leguminosae). PhytoKeys, 205, pp. 3. doi:10.3897/phytokeys.205.85866

Ringelberg, J. J., Koenen, E. J., Sauter, B., Aebli, A., Rando, J. G., Iganci, J. R., de Queiroz, L. P., Murphy, D. J., Gaudeul, M., Bruneau, A. & Luckow, M. (2023). Precipitation is the main axis of tropical plant phylogenetic turnover across space and time. Science Advances, 9(7), pp. eade4954. doi:10.1126/sciadv.ade4954

Rosseel, Y. (2012). *lavaan*: an R package for structural equation modelling. Journal of Statistical Software, 48, pp. 1–36. doi:10.18637/jss.v048.i02

Sandom, C., Faurby, S., Sandel, B. & Svenning, J.-C. (2014). Global late Quaternary megafauna extinctions linked to humans, not climate change. Proceedings of the Royal Society B, 281(1787), pp. 20133254. doi:10.1098/rspb.2013.325

Smith, F. A., Boyer, A. G., Brown, J. H., Costa, D. P., Dayan, T., Ernest, S. M., Evans, A. R., Fortelius, M., Gittleman, J. L., Hamilton, M. J. & Harding, L. E. (2010). The evolution of maximum body size of terrestrial mammals. Science, 330(6008), pp. 1216–1219. doi:10.1126/science.1194830

Staver, A. C., Bond, W. J., Cramer, M. D. & Wakeling, J. L. (2012). Top-down determinants of niche structure and adaptation among African Acacias. Ecology Letters, 15, pp. 673–679. doi:10.1111/j.1461-0248.2012.01780.x

Stuart, A. J. (2015). Late Quaternary megafaunal extinctions on the continents: a short review. Geological Journal, 50, pp. 338–363. doi:10.1002/gj.2710

Tomlinson, K. W., van Langevelde, F., Ward, D., Prins, H. H., de Bie, S., Vosman, B., Sampaio, E. V. & Sterck, F. J. (2016). Defence against vertebrate herbivores trades off into architectural and low nutrient strategies amongst savanna Fabaceae species. Oikos, 125(1), pp. 126–136. doi:10.1111/oik.02178

Valverde, P. L., Fornoni, J. & Núñez-Farfán, J. (2001). Defensive role of leaf trichomes in resistance to herbivorous insects in *Datura stramonium*. Journal of Evolutionary Biology, 14, pp. 424–432. doi:10.1046/j.1420-9101.2001.00307.x

Van Langevelde, F., Van De Vijver, C. A., Kumar, L., Van De Koppel, J., De Ridder, N., Van Andel, J., Skidmore, A. K., Hearne, J. W., Stroosnijder, L., Bond, W. J. & Prins, H. H. (2003). Effects of fire and herbivory on the stability of savanna ecosystems. Ecology, 84(2), pp. 337–350. doi: 10.1890/0012-9658(2003)084[0337:EOF AHO]2.0.CO;2

War, A. R., Paulraj, M. G., Ahmad, T., Buhroo, A. A., Hussain, B., Ignacimuthu, S. & Sharma, H. C. (2012). Mechanisms of plant defense against insect herbivores. Plant Signaling & Behavior, 7, pp. 1306–1320. doi:10.4161/psb.21663

Wölke, F. J. R., Onstein, R. E., Kissling, W. D. & Linder, H. P. (2023). Africa as an evolutionary arena for large fruits. New Phytologist, 240(4), pp. 1574–1586. doi:10.1111/nph.19456

Zhang, X., Gélin, U., Spicer, R. A., Wu, F., Farnsworth, A., Chen, P., Del Rio, C., Li, S., Liu, J., Huang, J. & Spicer, T. E. (2022). Rapid Eocene diversification of spiny plants in subtropical woodlands of central Tibet. Nature Communications, 13(1), pp. 3787. doi:10.1038/s41467-022-31211-3

