## Supplementary Material for "The complex interaction between megaherbivores, climate and fire has shaped the evolution and distribution of plant spinescence across biogeographical realms"

**Institutional affiliations:**

**Figure S1: Effect of minimum number of mimosoids per grid cell on global structural equation modelling (SEM) results.** Structural equation models showing the direct and indirect effects of extant and extinct megaherbivore species richness and environmental factors on the proportion of spinescent mimosoid species across assemblages with (a) a minimum of 3 mimosoid species (N = 2192 grid cells) (also see main text), (b) a minimum of 1 mimosoid species (N = 3156 grid cells) and (c) a minimum of 5 mimosoid species (N = 1649 grid cells).

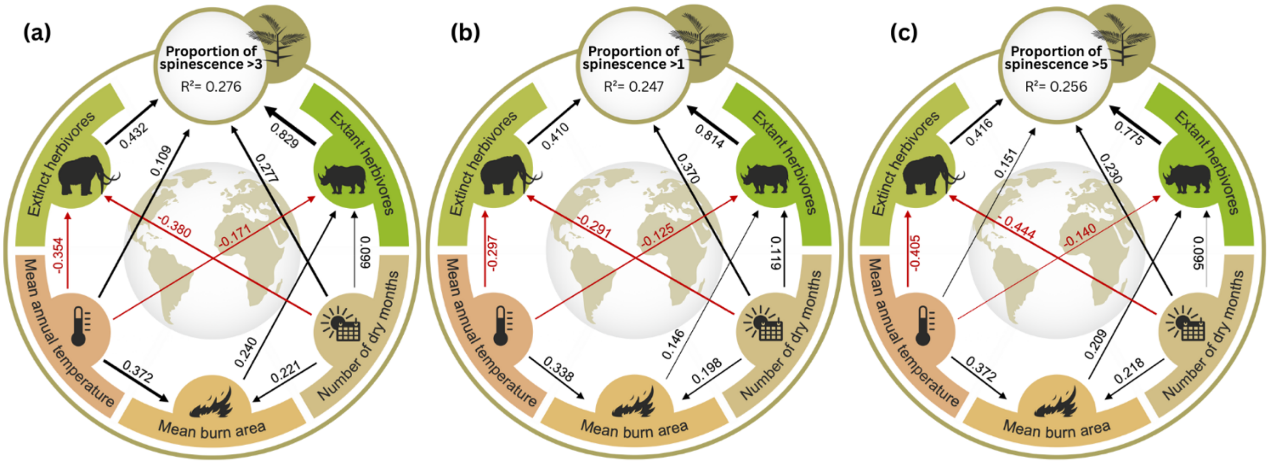

**Figure S2: Effect of extant and extinct herbivore richness, length of the driest season and mean annual temperature on randomised global assemblages with spinescence**. The standardized coefficients on the proportion of spinescent species from the observed/empirical model (red line) as derived from SEMs, is compared to 1,000 simulated effects (frequency distributions in grey). These simulations were conducted by randomly assigning the presence or absence of defense traits (spinescence) across mimosoid species, while preserving the total species richness of mimosoids for each grid cell. The observed effect of both extant and extinct herbivore richness, were higher than expected from a random distribution, and most of the effect for length of the driest season and mean annual temperature on the proportion of spinescent species, were also higher than expected from a random distribution of spinescence.

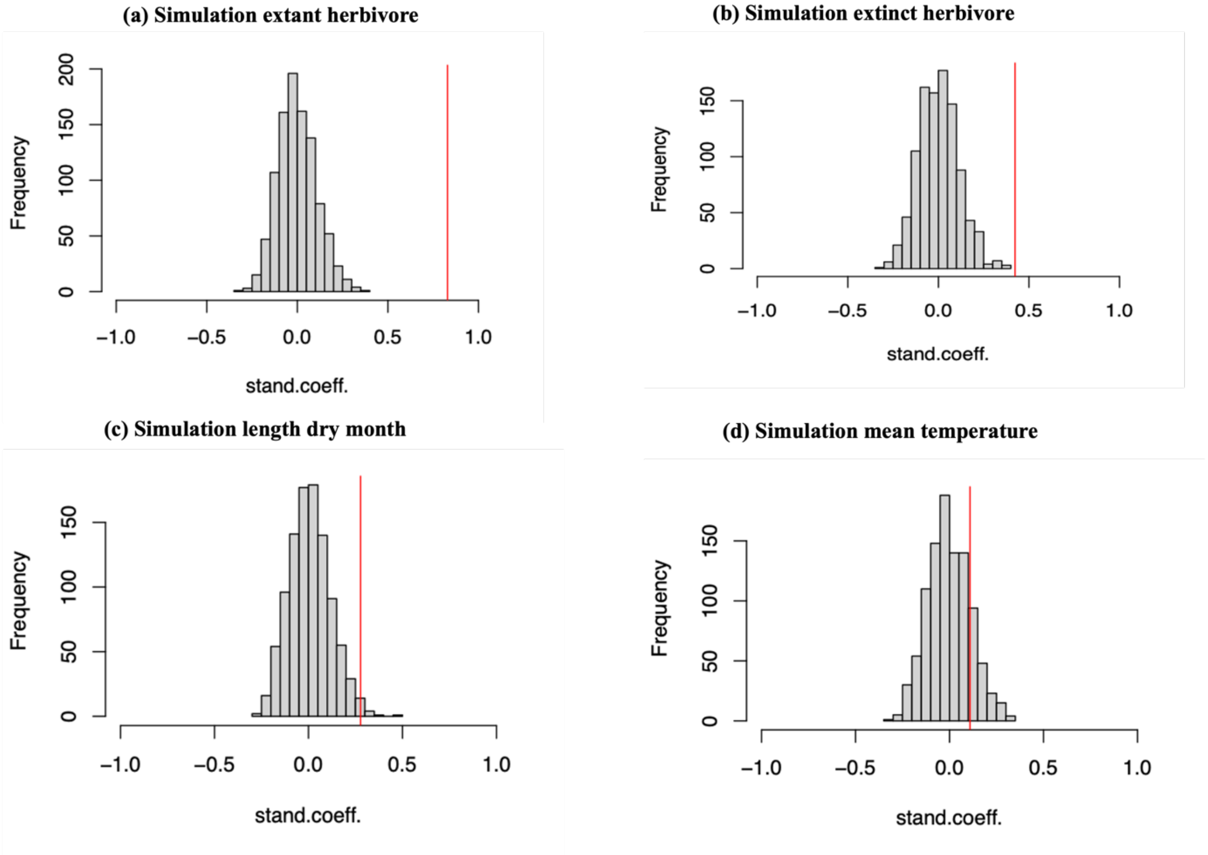

**Table S1: Glossary of morphological armature states in Mimoseae.** This table summarizes six categories of physical defense structures.

| **Armature state** | **Description** | **Origin** |
|---|---|---|
| Unarmed | No prickles, spines or thorns present | None |
| Stipular spines | Leaf-base stipules modified into rigid, pointed spines | Derived from modified stipules at the petiole base |
| Spinescent shoots | Entire lateral shoots or shoot segments transformed into stiff, spine-like structures | Shoots (including short shoots) becoming spine-like |
| Thorns | Woody, branch-derived thorns formed from axillary bud | Modified axillary branch |
| Prickles | Sharp epidermal/cortical outgrowths along node or internode lacking vascular tissue | Stem nodes |
| Axillary spine | Single sharp structures emerging from the leaf axil, distinct from stipular or branch thorns | Modified axillary bud scale or bud trace |

**Table S2a: Standardized regression coefficients from the global structural equation model (SEM) analyzing predictors of spinescence in mimosoid legumes**. Predictor variables include extant and extinct herbivore richness, dry season length, mean annual temperature (*temp_chelsa*), and fire frequency (burned area; BA). Coefficients reflect direct paths only. All models were estimated using maximum likelihood with robust standard errors.

**Global SEM**

| **Statistic** | **Value** |
|---|---|
| Estimator | ML |
| Optimization method | NLMINB |
| Number of parameters | 17 |
| Observations (used / total) | 2192 / 2685 |
| Chi-Square (df = 1, p) | 2.704 (1), 0.100 |
| CFI | 0.999 |
| TLI | 0.987 |
| RMSEA | 0.028 |
| • 90 % CI | 0.000 – 0.070 |
| • p (RMSEA ≤ 0.05) | 0.757 |
| • p (RMSEA ≥ 0.08) | 0.018 |
| SRMR | 0.007 |
| AIC | –2705.464 |
| BIC | –2608.690 |
| SABIC | –2662.702 |

| **Outcome** | **Predictor** | **Estimate** | **Std.Err** | **z-value** | **p-value** | **Std.lv** | **Std.all** |
|---|---|---|---|---|---|---|---|
| perc_spines | extant | 0.829 | 0.032 | 26.125 | 0.000 | 0.829 | 0.514 |
| | extinct | 0.423 | 0.022 | 19.396 | 0.000 | 0.423 | 0.387 |
| | lengthDry | 0.277 | 0.027 | 10.150 | 0.000 | 0.277 | 0.195 |
| | temp_chelsa | 0.109 | 0.045 | 2.439 | 0.015 | 0.109 | 0.046 |
| extant | temp_chelsa | –0.171 | 0.032 | –5.321 | 0.000 | –0.171 | –0.116 |
| | lengthDry | 0.099 | 0.019 | 5.183 | 0.000 | 0.099 | 0.113 |
| | BA | 0.240 | 0.025 | 9.588 | 0.000 | 0.240 | 0.203 |
| extinct | temp_chelsa | –0.354 | 0.045 | –7.919 | 0.000 | –0.354 | –0.162 |
| | lengthDry | –0.380 | 0.027 | –14.269 | 0.000 | –0.380 | –0.292 |
| BA | temp_chelsa | 0.372 | 0.025 | 15.039 | 0.000 | 0.372 | 0.298 |
| | lengthDry | 0.221 | 0.015 | 15.023 | 0.000 | 0.221 | 0.298 |

| **R- Square** | **Estimate** |
|---|---|
| Perc_spine | 0.276 |
| extant | 0.070 |
| extinct | 0.099 |
| BA | 0.154 |

**Table S2b: Standardized coefficients from the global Spatial Autoregressive Model (SAR) model.** This model included the same predictors on the proportion of spinescent mimosoids as in Table S1a. The model accounts for spatial autocorrelation using a spatial weights matrix based on geographic proximity. Global SAR Error Model (N = 2685).

| Predictor | Estimate | Std. Error | z-value | p-value |
|---|---|---|---|---|
| (Intercept) | –0.2143 | 0.0529 | –4.0496 | $5.13 \times 10^{-5}$ |
| extant | 0.7858 | 0.0288 | 27.2456 | $< 2.2 \times 10^{-16}$ |
| extinct | 0.4484 | 0.0193 | 23.2202 | $< 2.2 \times 10^{-16}$ |
| lengthDry | 0.3385 | 0.0203 | 16.6957 | $< 2.2 \times 10^{-16}$ |
| temp_chelsa | 0.0863 | 0.0424 | 2.0336 | $4.20 \times 10^{-2}$ |

**Table S3a: Standardized regression coefficients from the structural equation model (SEM) applied to the Americas subset.** The model includes direct and indirect effects of environmental and biotic predictors on the proportion of spinescent species. All pathways and model fit statistics follow the same structure as the global SEM (Table S1a).

| Statistic | Value |
|---|---|
| Estimator | ML |
| Optimization method | NLMINB |
| Number of parameters | 15 |
| Observations (used / total) | 1036 / 1282 |
| Chi-Square (df = 2, p) | 6.122 (3), 0.106 |
| CFI | 0.996 |
| TLI | 0.984 |
| RMSEA | 0.032 |
| • 90 % CI | 0.052 – 0.126 |
| • p (RMSEA ≤ 0.05) | 0.759 |
| • p (RMSEA ≥ 0.08) | 0.011 |
| SRMR | 0.016 |
| AIC | – 5026.320 |
| BIC | – 4952.173 |
| SABIC | – 4999.815 |

| Outcome | Predictor | Estimate | Std.Err | z-value | p-value | Std.lv | Std.all |
|---|---|---|---|---|---|---|---|
| perc_spines | extant | 0.829 | 0.166 | 5.010 | 0.000 | 0.829 | 0.140 |
| | extinct | 0.125 | 0.029 | 4.275 | 0.000 | 0.125 | 0.115 |
| | lengthDry | 0.648 | 0.033 | 19.736 | 0.000 | 0.648 | 0.498 |
| | temp_chelsa | –0.477 | 0.055 | –8.679 | 0.000 | –0.477 | –0.222 |
| extant | lengthDry | 0.031 | 0.006 | 5.082 | 0.000 | 0.031 | 0.141 |
| | temp_chelsa | –0.070 | 0.010 | –6.814 | 0.000 | –0.070 | –0.193 |
| | BA | –0.056 | 0.009 | –5.970 | 0.000 | –0.056 | –0.179 |
| extinct | BA | 0.640 | 0.049 | 12.934 | 0.000 | 0.640 | 0.373 |
| BA | temp_chelsa | 0.221 | 0.035 | 6.272 | 0.000 | 0.221 | 0.191 |

| R- Square | Estimate |
|---|---|
| Perc_spine | 0.358 |
| extant | 0.109 |
| extinct | 0.145 |
| BA | 0.037 |

**Table S3b. Standardized coefficients from the global Spatial Autoregressive Model (SAR) model in the Americas.** This model included the same predictors on the proportion of spinescent mimosoids as in Table S1a. The model accounts for spatial autocorrelation using a spatial weights matrix based on geographic proximity.

| **Predictor** | **Estimate** | **Std. Error** | **z-value** | **p-value** |
|---|---|---|---|---|
| (Intercept) | 0.1873 | 0.0481 | 3.8906 | $1.00 \times 10^{-4}$ |
| extant | 0.5478 | 0.1704 | 3.2156 | $1.30 \times 10^{-3}$ |
| extinct | 0.0878 | 0.0301 | 2.9118 | $3.59 \times 10^{-3}$ |
| lengthDry | 0.5124 | 0.0260 | 19.7097 | $< 2.2 \times 10^{-16}$ |
| temp_chelsa | –0.2874 | 0.0543 | –5.2919 | $1.21 \times 10^{-7}$ |

**Table S4a: Standardized regression coefficients from the structural equation model (SEM) applied to the Africa subset.** The model includes direct and indirect effects of environmental and biotic predictors on the proportion of spinescent species. All pathways and model fit statistics follow the same structure as the global SEM (Table S1a).

| **Statistic** | **Value** |
|---|---|
| Estimator | ML |
| Optimization method | NLMINB |
| Number of parameters | 16 |
| Observations (used / total) | 443 / 511 |
| Chi-Square (df = 2, p) | 1.418 (2), 0.492 |
| CFI | 1.000 |
| TLI | 1.005 |
| RMSEA | 0.000 |
| • 90 % CI | 0.000 – 0.085 |
| • p (RMSEA ≤ 0.05) | 0.766 |
| • p (RMSEA ≥ 0.08) | 0.066 |
| SRMR | 0.012 |
| AIC | –2077.890 |
| BIC | –2012.393 |
| SABIC | –2063.170 |

| **Outcome** | **Predictor** | **Estimate** | **Std.Err** | **z-value** | **p-value** | **Std.lv** | **Std.all** |
|---|---|---|---|---|---|---|---|
| perc_spines | extinct | 1.389 | 0.231 | 6.004 | 0.000 | 1.389 | 0.282 |
| | extant | 0.245 | 0.072 | 3.424 | 0.001 | 0.245 | 0.146 |
| | lengthDry | 1.030 | 0.105 | 9.846 | 0.000 | 1.030 | 0.400 |
| extant | lengthDry | 0.211 | 0.064 | 3.305 | 0.001 | 0.211 | 0.137 |
| | temp_chelsa | –0.849 | 0.080 | –10.776 | 0.000 | –0.849 | –0.444 |
| | BA | 0.310 | 0.057 | 5.480 | 0.000 | 0.310 | 0.228 |
| extinct | lengthDry | 0.213 | 0.018 | 11.971 | 0.000 | 0.213 | 0.406 |
| | temp_chelsa | –0.347 | 0.022 | –15.478 | 0.000 | –0.347 | –0.532 |
| | BA | 0.057 | 0.016 | 3.593 | 0.000 | 0.057 | 0.122 |
| BA | temp_chelsa | 0.226 | 0.065 | 3.497 | 0.001 | 0.226 | 0.161 |

| **R- Square** | **Estimate** |
|---|---|
| Perc_spine | 0.411 |
| extant | 0.252 |
| extinct | 0.501 |
| BA | 0.026 |

**Table S4b: Standardized coefficients from the global Spatial Autoregressive Model (SAR) model in Africa.** This model included the same predictors on the proportion of spinescent mimosoids as in Table S1a. The model accounts for spatial autocorrelation using a spatial weights matrix based on geographic proximity.

| Predictor | Estimate | Std. Error | z-value | p-value |
|---|---|---|---|---|
| (Intercept) | –0.7659 | 0.0755 | –10.1506 | $< 2.2 \times 10^{-16}$ |
| extant | 0.2480 | 0.0707 | 3.5073 | $4.53 \times 10^{-4}$ |
| extinct | 1.5224 | 0.2185 | 6.9686 | $3.20 \times 10^{-12}$ |
| lengthDry | 1.3139 | 0.0941 | 13.9696 | $< 2.2 \times 10^{-16}$ |

**Table S5: Transition Rates for Different Armature Types under ER and ARD Models.** This table presents the transition rates for each armature type (Thorn, Spinescent shoot, Prickles, Stipular Spines, Axillary Spines, and Armed/Unarmed) under the Equal Rates (ER) and All Rates Different (ARD) models. Transition rates are reported for both directions between "Other" and the armature type. AIC values for each model are provided to indicate the fit of the respective models, with best-fit (ARD) models highlighted in **bold**.

| **Armature Type** | **Model** | **AIC** | ***Other* to Armature Rate** | **Armature to *Other* Rate** |
|---|---|---|---|---|
| Thorn | ER | 75.94366 | 0.0005495872 | 0.0005495872 |
| **Thorn** | **ARD** | **73.37522** | **0.0180358185** | **0.0004964227** |
| Spinescent shoot | ER | 179.8190 | 0.001293418 | 0.001293418 |
| **Spinescent shoot** | **ARD** | **159.3105** | **0.188780711** | **0.001996742** |
| Prickles | ER | 654.2586 | 0.006296967 | 0.006296967 |
| **Prickles** | **ARD** | **557.0988** | **0.0408567385** | **0.0008409105** |
| Stipular Spines | ER | 388.0017 | 0.003877855 | 0.003877855 |
| **Stipular Spines** | **ARD** | **386.6190** | **0.011122861** | **0.003548655** |
| Axillary Spines | ER | 35.53843 | 0.000182519 | 0.000182519 |
| **Axillary Spines** | **ARD** | **29.00333** | **0.0001503062** | **0.1324078788** |
| Armed/Unarmed | ER | 926.3572 | 0.01162165 | 0.01162165 |
| **Armed/Unarmed** | **ARD** | **909.6673** | **0.008185383** | **0.023150566** |
